## Supplementary figures and images for "The haemodynamics of the human placenta in utero"

### S1 Fig

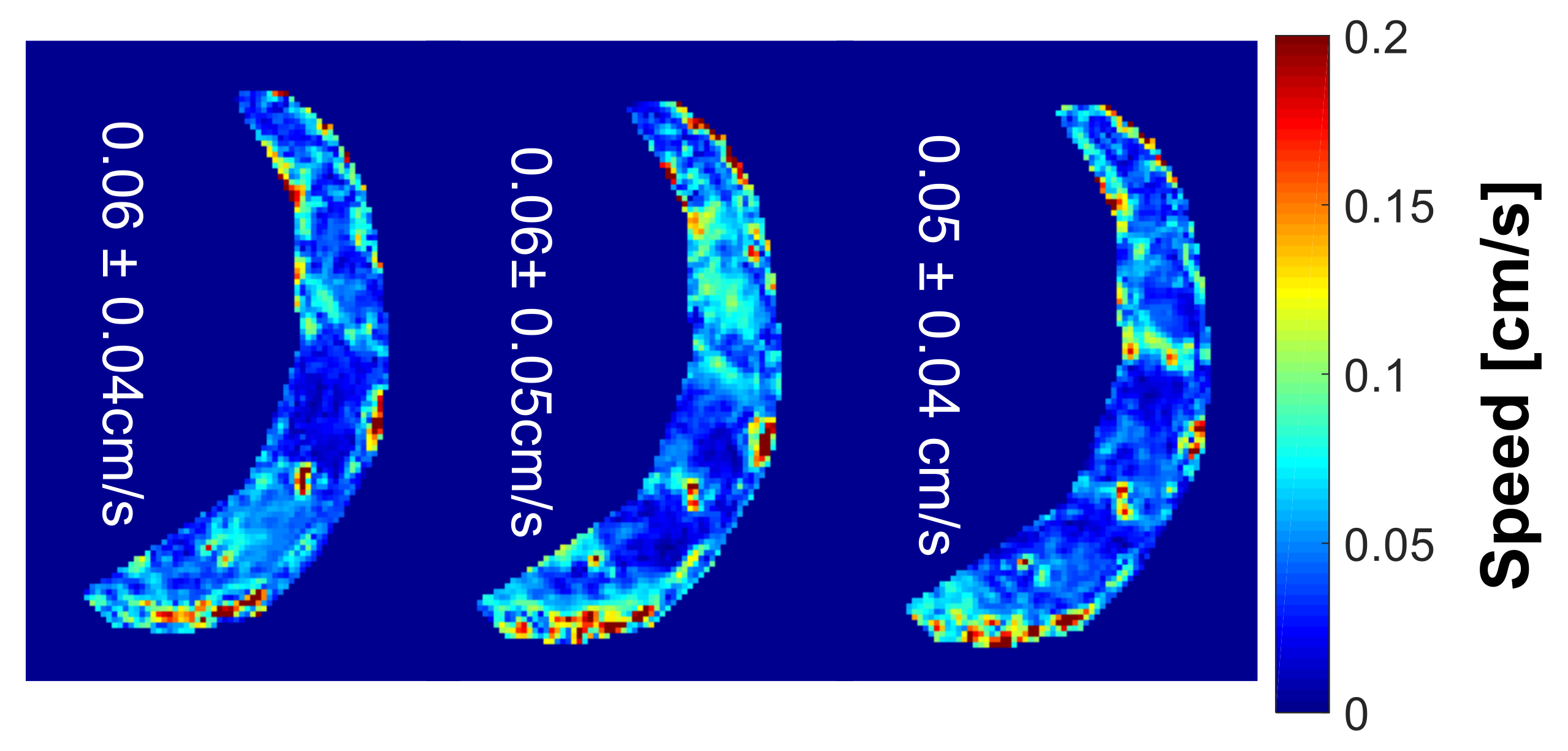

### S2 Fig

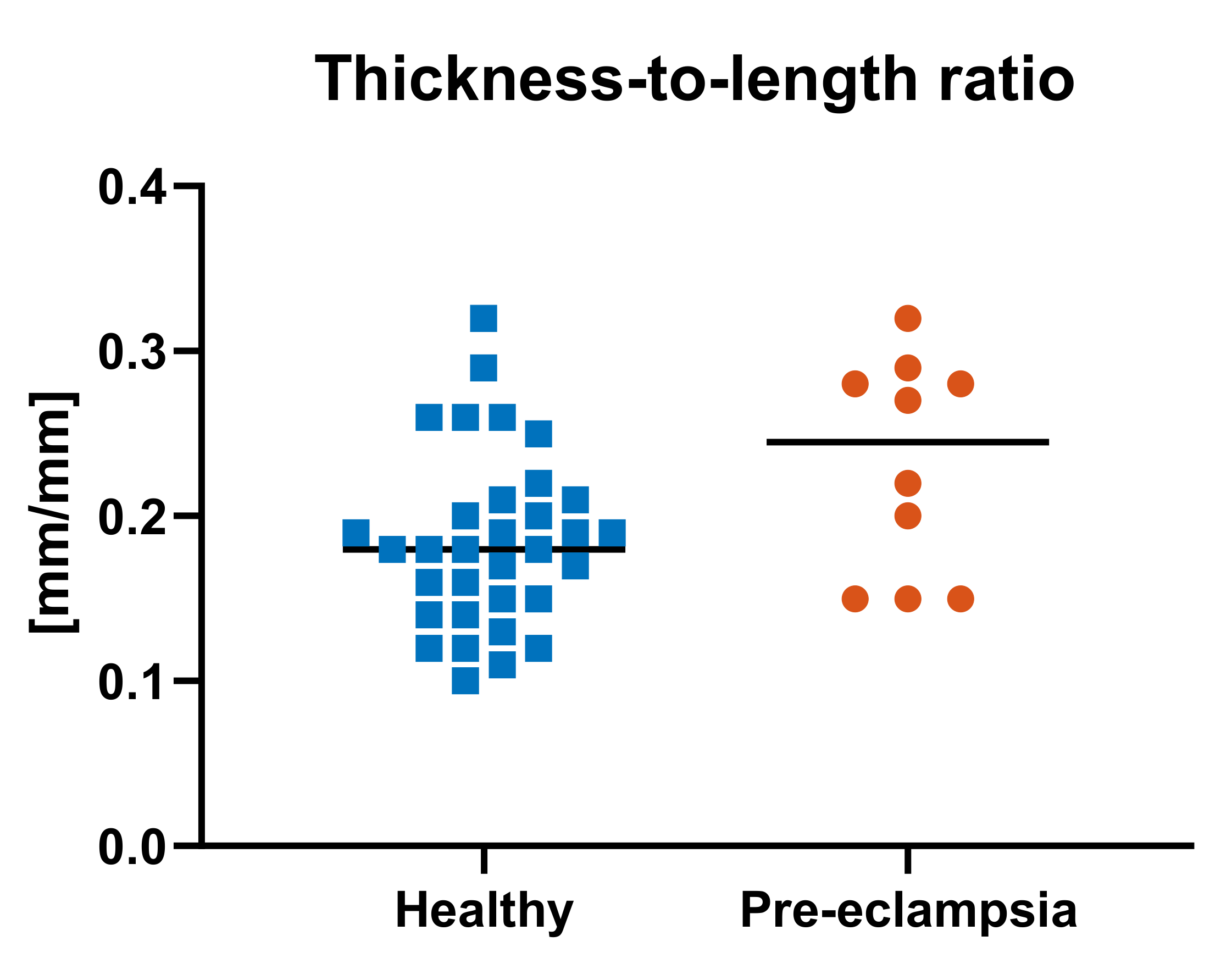

### S3 Fig

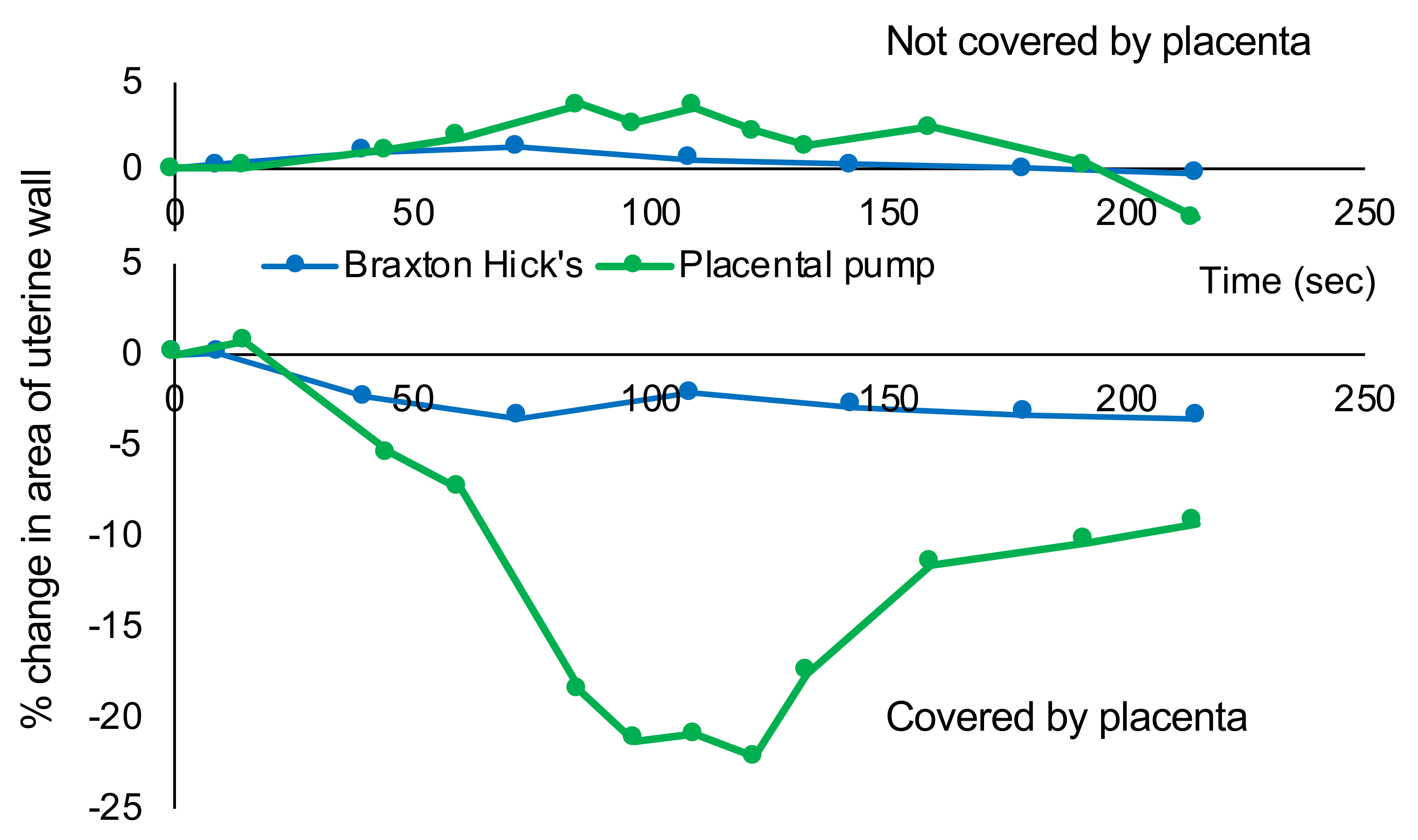

### S4 Fig

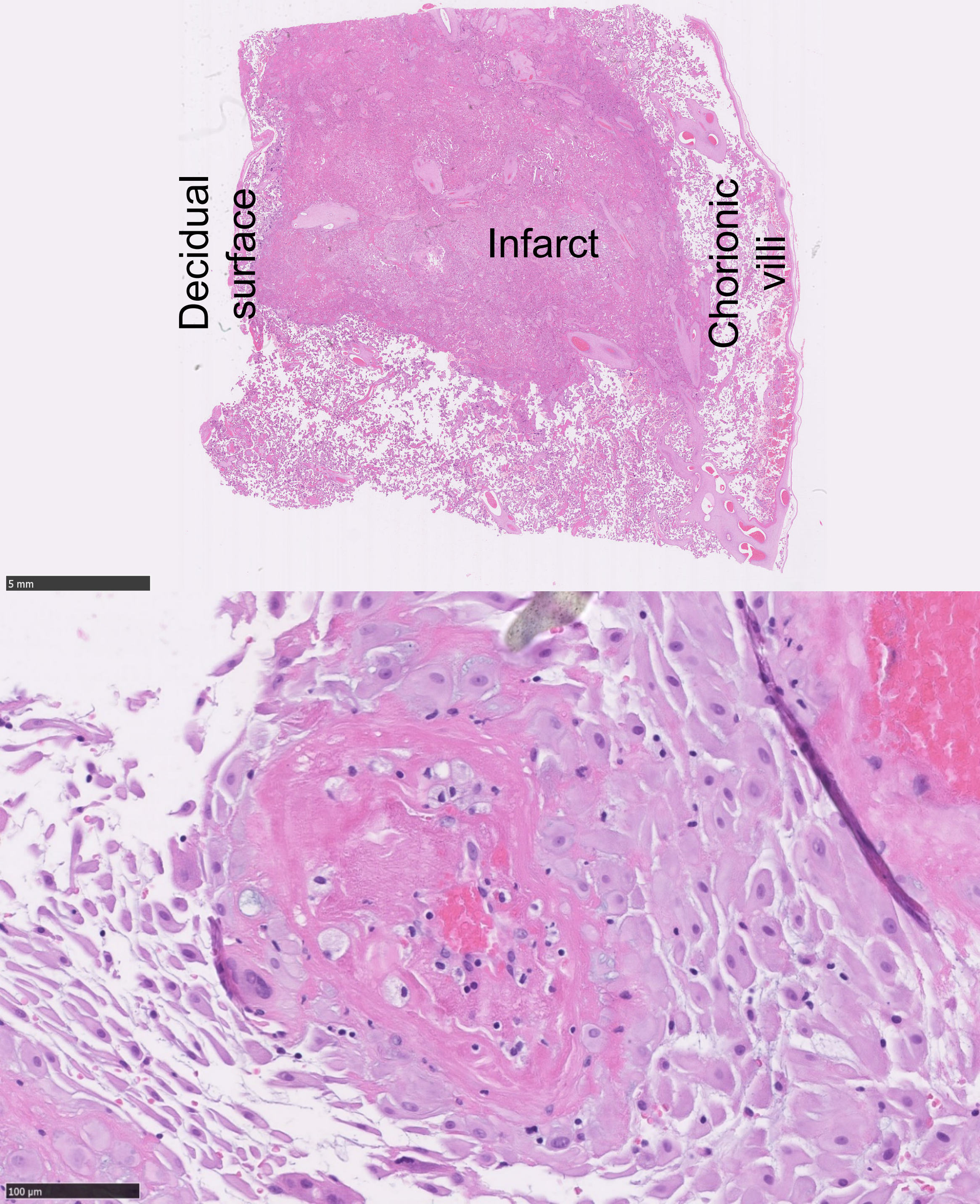

### S5 Fig

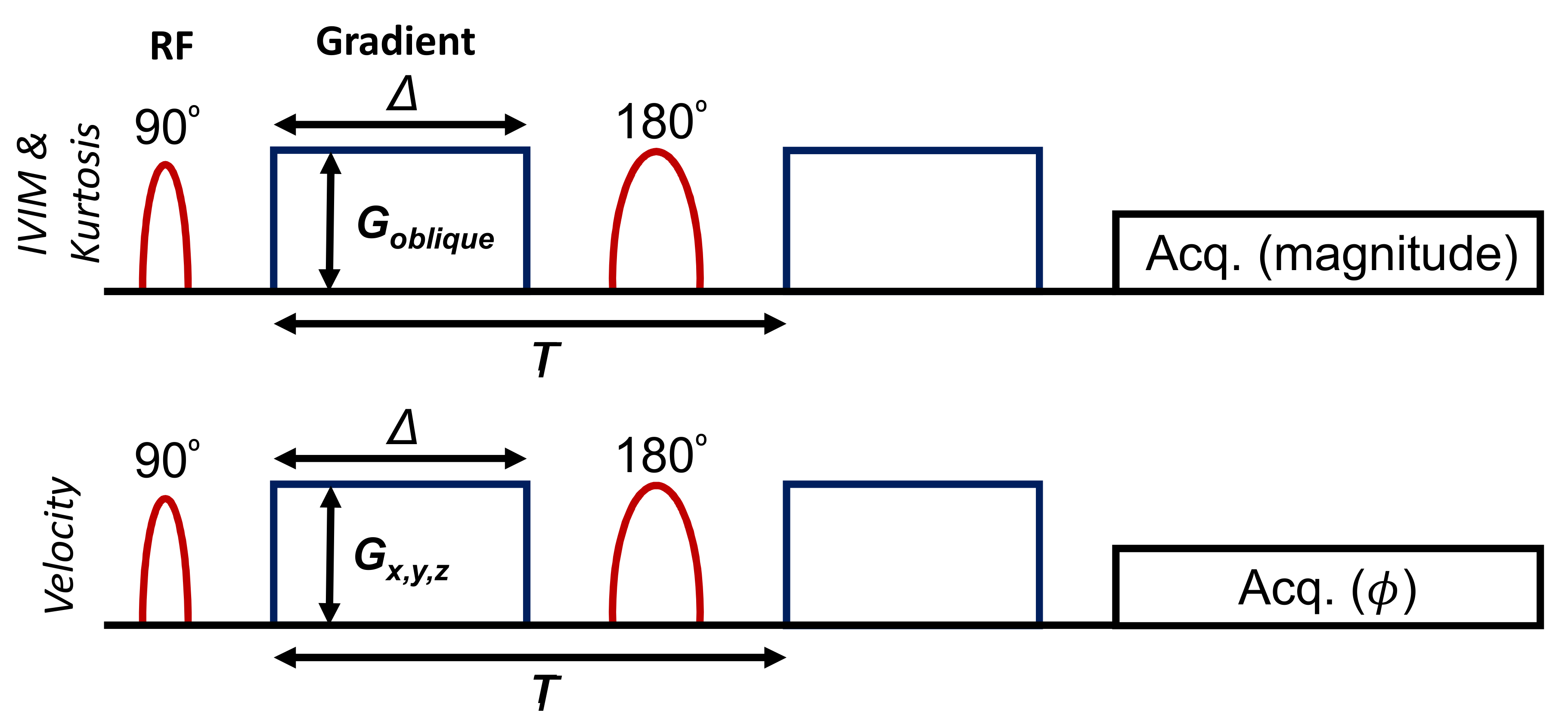

### S6 Fig

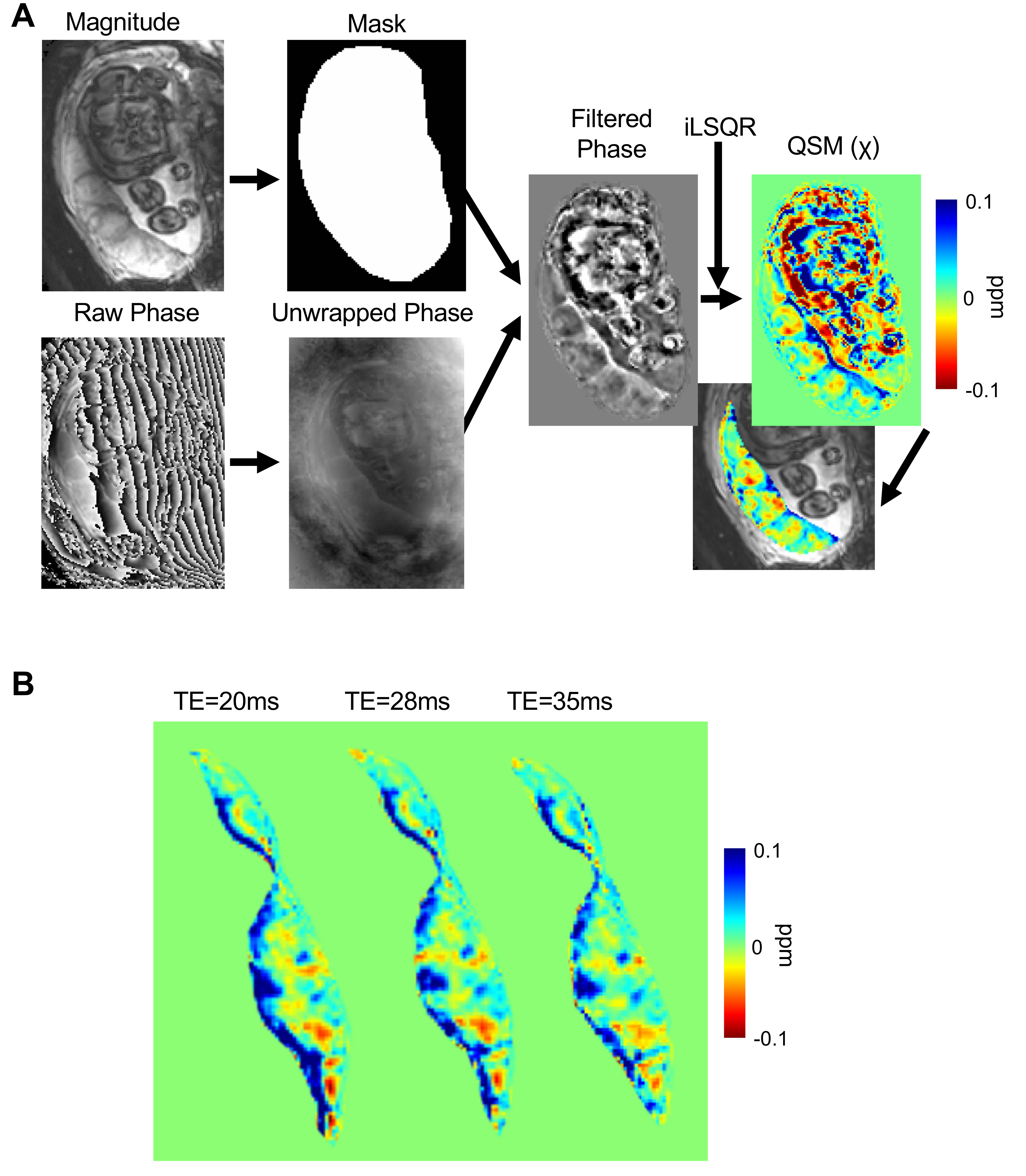
